# Identifying nearshore nursery habitats for sharks in the Eastern Tropical Pacific from fisheries landings and interviews

**DOI:** 10.1101/2021.02.03.429561

**Authors:** Juliana López-Angarita, Melany Villate, Juan Manuel Díaz, Juan C. Cubillos, Alexander Tilley

## Abstract

The Eastern Tropical Pacific (ETP) comprising the coasts of Costa Rica, Panama, Colombia and Ecuador, represents an area of high marine biodiversity that supports productive fisheries and acts as an important migratory corridor for many marine species. Despite its biological importance, the ETP is understudied and lacks sufficient data for science-based fisheries management and conservation decision-making. This study aims to consolidate understanding of the current and historical distribution of sharks and mobulid rays in the ETP. We used interviews of coastal community stakeholders to document traditional knowledge of shark and mobulid ray species and distributions. We also analysed small-scale fisheries landings data, where available, to quantify local exploitation patterns and the importance of sharks and rays in small-scale fisheries catches. All shark species landed in the dominant nearshore gillnet fishery show very low mean individuals weights (<5 kg), indicating that the fisheries are dominated by juveniles, captured. Aside from smooth-hounds (*Mustelus* spp.), the scalloped hammerhead shark, *Sphyrna lewini*, is the most frequently landed shark species in the region by weight and number, with peaks in abundance between April - July. From 132 interviews in 51 communities across the three countries, and landings data from two small-scale fisheries sites, we identified 41 sites in 12 broad geographical zones as important shark nursery habitats. Of these sites, 68% were associated closely with large mangrove systems of the ETP, highlighting the importance of this habitat for shark life history. No patterns were seen in the occurrence or distribution of mobulid rays in coastal areas. Marine protected areas and responsible fishing zones cover 37% of identified nursery habitats in the ETP, 30% in Costa Rica, 48% in Panama and 30% in Colombia. These findings provide an important benchmark of the conservation status of sharks in the ETP and allow for the prioritisation of research and policy-making.

## Introduction

The Eastern Tropical Pacific is a biologically rich region (Miloslavich *et al*., 2011) that encompasses the Pacific coasts of Panama, Costa Rica, Ecuador and Colombia. Its oceanographic characteristics and climate regimes make it a very dynamic and productive region (Fiedler and Lavín, 2006; Wang and Fiedler, 2006). Consequently, the ETP harbours great biodiversity and productivity of marine faunal biomass (Ryther, 1969; Fiedler, Philbrick and Chavez, 1991) and is a major corridor for migratory species (Nasar *et al*., 2016).

Sharks and mobulid rays are very vulnerable to fishing pressure because of life-history traits such as late maturation, slow growth, and limited fecundity (Fowler *et al*., 2005; Myers and Worm, 2005). Even though some species are migratory, with large home ranges, most rely on coastal ecosystems as feeding grounds, and during early and reproductive life stages (Heupel, Carlson and Simpfendorfer, 2007). There is substantial evidence that elasmobranchs rely on habitat near the continental shelf, with neonates and juveniles showing usage of core nursery grounds and site-attached coastal movements (Stevens *et al*., 2005; Garla *et al*., 2006; Kinney and Simpfendorfer, 2009). Recent studies have brought to light the unsustainable level of fishing pressure on shark and ray populations in the ETP (Chávez, 2007; Clarke *et al*., 2018; Zanella, López-Garro and Cure, 2019).

Fisheries in the ETP comprise artisanal, semi-industrial and industrial boats that target commercial species (e.g. mahi-mahi, tuna, jacks, weakfish, Pacific bearded brotula) with a considerable amount of incidental catch of non-targeted species, and underreported overall catches (Harper *et al*., 2014). A large proportion of the bycatch is sharks and rays caught on long-lines or trapped in gill nets of fishing vessels (Dapp *et al*., 2013; Harper *et al*., 2014; Queiroz *et* al., 2016; Griffiths, Kesner-Reyes and Garilao, 2019). Despite international and national legislation to halt shark-finning, retention and landing of several shark species^1^, other socio-economic factors such as finding alternative fisheries resources, the increasing demand and rise in the price of elasmobranchs’ fins and meat, play an important role in unreported catches (Dulvy *et al*., 2014). The ETP countries are steadily strengthening their protection of sharks and rays by incorporating them in national fisheries management plans and taking part in global initiatives such as the International Plan of Action for Conservation and Management of Sharks^2^. Since early 2000, nations have voluntarily drafted and adopted such country plans and just in the last 15 years have been implemented in the countries covering the ETP. Nonetheless, implementation at a regional level for the ETP is still pending (RPOA-Sharks, FAO, 2020). However, as policy changes are slow due to limited regional scientific capacity leading to a lack of scientific information upon which to base decisions (López-Angarita *et al*., 2014; Saavedra-Díaz, Pomeroy and Rosenberg, 2016).

A total of 176 species of sharks and rays are registered as occurring in the ETP (88 sharks and 88 rays) in 66 genera (34 shark and 32 ray genera) and 33 families (19 and 14, respectively). Previous work on elasmobranch population ecology in the ETP has traditionally focused on adult pelagic shark populations (such as *Sphyrna lewini*) around the iconic oceanic islands of Galapagos, Cocos, and Malpelo (Bessudo *et al*., 2011; Ketchum *et al*., 2014; Hearn *et al*., 2016; Nalesso *et* al., 2019). There is convincing evidence of the genetic connectivity between these ETP oceanic islands and the habitats of the continental shelf (Quintanilla et al. (2015), and a current initiative is investigating the Gulf of Tribuga as nursery habitat for hammerhead sharks tagged in Malpelo Island (Bessudo, Ladino and Camargo, 2018). Little attention has been focused on elasmobranch use of coastal areas, or on the drivers of change from fishing communities whose livelihood activities are interwoven with these marine resources. However, recent research in Costa Rica has provided more insight into the populations of sharks and rays in the country (Espinoza *et al*., 2018, 2020).

The coastline of the ETP is largely covered by mangrove forests, that are known to support myriad fish and shark species (Edgar *et al*., 2011; Llerena *et al*., 2015; Espinoza *et al*., 2020), as well as providing diverse ecosystem services and resources that support livelihoods and food security for coastal communities (López-Angarita *et al*., 2016). The removal or alteration of coastal habitat combined with unsustainable fishing practices can accelerate the decline of these populations towards local extirpation (Harper *et al*., 2014). The rapid decline of elasmobranchs worldwide increases the urgency with which local ecological knowledge and scientific analysis of critical habitats to shark and ray populations must be recorded and generated.

Information on the presence, abundance, and distribution of critical habitat of sharks in the ETP is required to identify priorities and streamline policy advice around the sustainable management of shark populations as part of fisheries management at national and regional levels. In this study, we aim to use small-scale fisheries landings data and fishery stakeholder interviews to elucidate the number and locations of potential elasmobranch critical habitats along the Pacific coasts of Colombia, Panama and Costa Rica. We then compare these data with existing protection and management measures offered by the regional marine protected area network, seasonal closures, and selective fishing regulations on gears and species across the study countries.

## Methods

### Study sites

This study took place on the Pacific coasts of Costa Rica (CR), Panama (PAN) and Colombia (COL) in the ETP region (Figure 1). Interviews were undertaken in Costa Rica in May 2013 and February 2014; in Panama between July and October 2014 and between August and October 2015; and in Colombia in September and October 2015. Interview sampling was designed to cover the entire coast of the three studied countries, however, the southern coast of Colombia was not sampled due to practical and logistical constraints.

**Figure 1:**
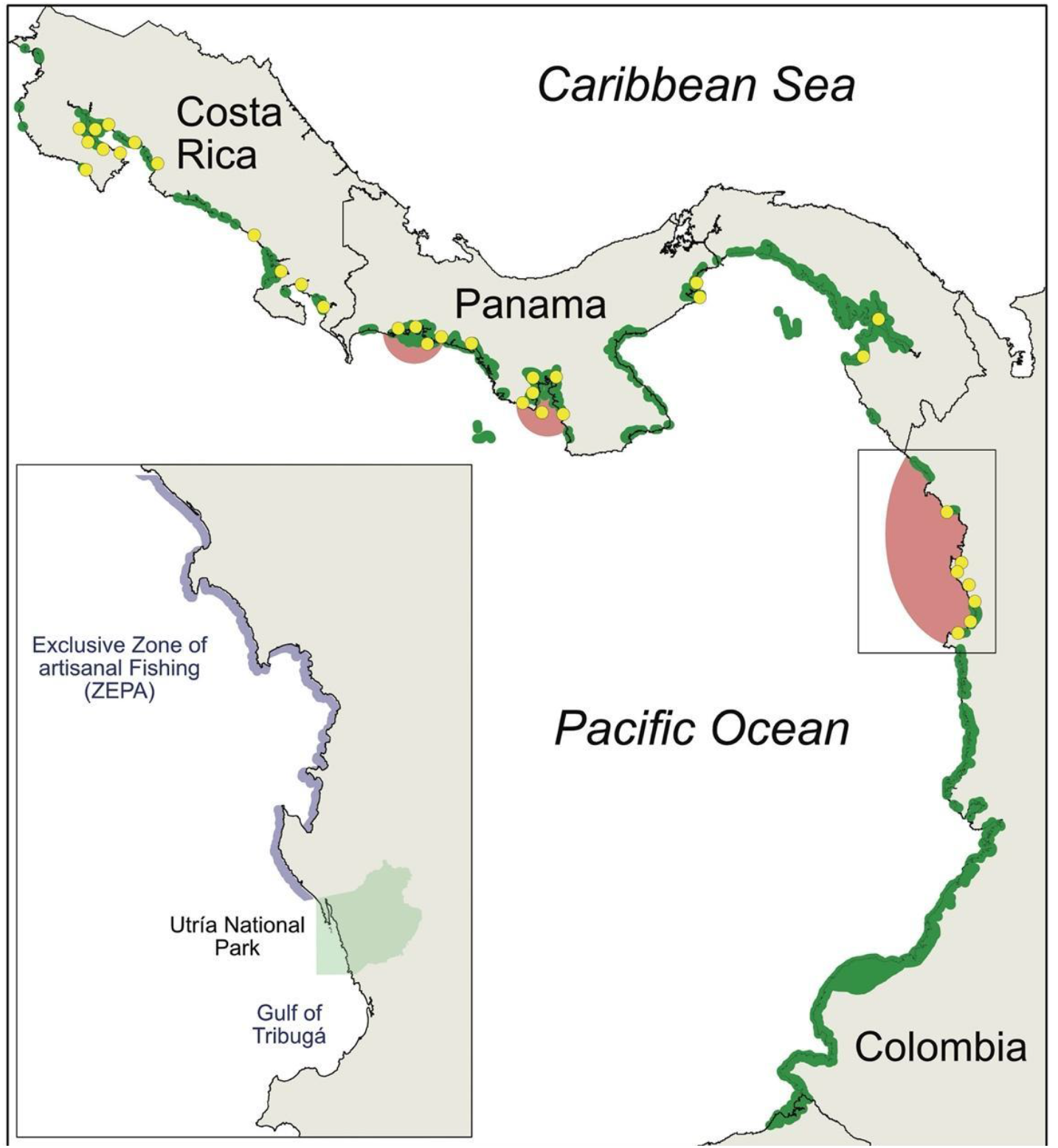
Map of study sites on the Eastern Tropical Pacific coasts of Costa Rica, Panama and Colombia with mangrove coverage (green coastal cord). Yellow points are locations of interviews. Red shading illustrates areas from which landings data were obtained and analysed. Inset map of Northern Chocó region of Colombia with different management zones. The Regional District of Integrated Management of the Gulf of Tribugá, Cabo Corrientes (DRMI) (60,138.6 ha) established in 2015, is not shown.

### Fisher Interviews

Semi-structured interviews were conducted in person by enumerators with fisheries stakeholders (fishers, traders, fishing cooperative managers, and community leaders) in 52 fishing communities (CR N=14, PAN N=25, COL N=7). A snowball sampling approach was used to identify respondents. The first point of contact was the manager or representative of a community fishing association (*coope*), and then these respondents provided referrals of others within their communities according to our area of inquiry. Given our interest in changing fisheries dynamics over time, one criterion for fisher respondents was that they had been fishing for at least 15 years. Interview questions were designed to gather information on: i) current and historical fishing activities, gears and catch trends, ii) past and current presence of sharks, rays and sawfish in the area iii) potential nursery grounds and perceived health of coastal habitats, and iv) perceived causes of observed environmental and fisheries trends (Appendix 1). Image cards of different elasmobranchs were used to assess fishers’ ability to identify different species, to record local names in use, and to prompt insights or records of catches and locations for the species. Printed maps of the local area were used to allow respondents to identify specific areas of interest if they were too distant to take enumerators to them in person.

**Image 1.**
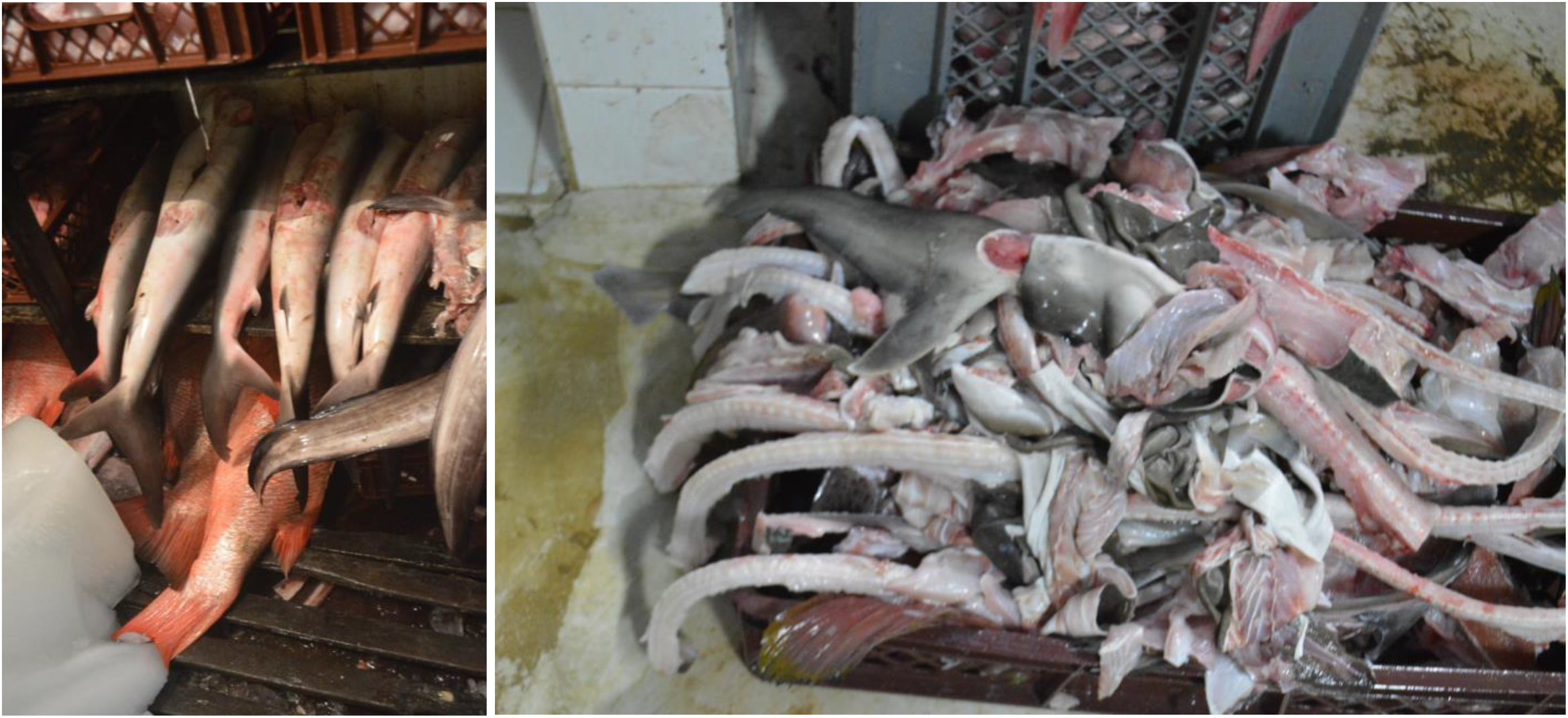
Juveniles sharks landed in Bahia Solano, Northern Choco, Colombia. © Juan Cubillos.

Initial fisher interviews and responses to ID cards suggested mobulid sightings were sporadic in the areas surveyed, so an additional questionnaire was designed to gather data from dive operators about possible mobulid aggregations exploited by tourism (Appendix 2). Surveys were conducted in situ or via telephone with dive shops in Costa Rica and Panama. This aspect was not carried out in Colombia due to very few dive operators in the area at the time of sampling. Interviews were recorded, transcribed and coded according to key themes. Qualitative analysis was performed using Nvivo software (version 11.1.1).

### Community site mapping

During interviews, respondents were asked if they were aware of, or could identify, particular areas of their fishing range where juvenile sharks were regularly sighted or caught, both currently and historically. Any areas mentioned were located by the respondent pointing to the location on a map, accompanying the interviewers to the site, or describing in as much detail as possible, the location and topography of the site. Geographical coordinates were taken, and where possible, areas identified by respondents were classified by species. When feasible, sites were verified with other respondents, and sites that were mentioned repeatedly by different respondents were ground-truthed with field visits. These ground-truthed locations in Costa Rica were: Isla Chira, Golfito, Playa Tortuga and Puerto Jiménez; Bahia Solano and Jurubidá in Colombia; and Pito, Bocachica and Cebaco in Panama. Enumerators accompanied fishing trips and recorded at landing sites to gather photos of individuals captured for identification purposes, along with information on the number, length and weight.

### Qualitative analysis

To determine the representation of critical habitat of elasmobranch species in the current MPA network of the ETP, we determined the percentage of points of identified critical habitat by interview respondents that were located inside or outside protected areas. We plotted the limits of marine reserves of Colombia, Costa Rica and Panama using the online interface of the World Database on Protected Areas, Protected Planet (www.protectedplanet.com). Given that fishers frequently identified mangrove habitats as nursery grounds for marine species, we overlaid the limits of protected areas with the global mangrove coverage dataset from Giri et al. (2011) available from the World Conservation Monitoring Centre (WCMC), and calculated the percentage of mangroves inside and outside protected areas using ArcGIS ver. 10.3 (ESRI).

### Small-scale fisheries landings

Through partnerships with local NGOs working in the region, we collated existing small-scale fisheries landings data sets. We restricted sampling to datasets that included a consistent measure of fishing effort, and that held more than one continuous year or data. No such dataset was possible to obtain for nearshore areas of Costa Rica at the time.

Datasets for SSF in Panama and Colombia were the result of a project implemented by the Marviva Foundation to train community members to collect fishery information using surveys designed for the Colombian national information system of landings (Neira *et al*., 2016). In Colombia, data were collected between March 2010 and September 2013 from 16 coastal communities in the Northern Choco region (Figure 1). In Panama, data were collected from 2012-2014 in one community in the Gulf of Montijo and two communities in the Gulf of Chiriqui. The information extracted from the dataset was catch volume and number of individuals of each species landed, number of fishers, trip duration (in days), fishing gear, and the landing site. Catch and effort data were recorded 3-4 times per week throughout the year, disrupted only by local religious holidays or inclement weather.

Species assemblages and relative importance of shark and ray species were summarised using an Index of Relative Importance (IRI), calculated as the combined proportion of total catch weight, frequency and number of individuals by location (Kolding, 1999). Catch per unit effort (CPUE) of sharks and rays using both kg/fisher*day and N/fisher*day, was compared between species, locations and over time. Analyses were performed in PASGEAR II ver. 2.8 (Kolding & Skaalevik) and JMP v.12 (SAS Inst.).

## Results

### Fisher interviews

In total, 132 interviews were performed across seven broad regions of Colombia, Panama, and Costa Rica (Table 1 and Figure 1). Respondents were predominantly men (124) and eight were women. The mean number of years of experience fishing across all respondents was 36 years. The youngest respondent was 25 years old, and the oldest 75 years (Figure 1).

**Table 1.** The number of interviews conducted by country and region.

| Country | Region | Number of communities | Number of interviews |
| --- | --- | --- | --- |
| Panama | Gulf of Chiriqui | 5 | 11 |
|  | Gulf of Montijo | 12 | 27 |
|  | Darien | 4 | 10 |
|  | West Panama City | 4 | 4 |
| Colombia | Northern Choco | 13 | 54 |
| Costa Rica | Gulf of Nicoya | 11 | 21 |
|  | Terraba Sierpe | 2 | 3 |

According to interviews in Colombia, juvenile sharks are caught using longlines, nets and hand lines usually close to the shore, using small paddle boats or motorized boats less than 8 m with low power engines. Once landed, they are used or sold as bait or for local consumption. Small sharks have a very low price in the market of ∼USD$0.50/kg. Some are sent to big cities such as Buenaventura, where they are intentionally mislabelled and sold as snook (*Centropomus undecimalis*). In Panama, juvenile sharks are mainly caught using gillnets in coastal areas and estuaries and are sold for ceviche (a locally popular dish of raw fish marinated in lemon juice). Interestingly, ceviche sellers in Panama city reported that they actively sought shark meat for their restaurants as it had better shelf-life; disintegrating slower compared to the weakfish (*Cynoscion spp*.) it was labelled as.

In Colombia, 94% of respondents mentioned that hammerhead sharks (*Sphyrna* spp.) were commonly found in their areas, and in Panama, this figure was 87%. Other species found in both countries were the blacktip shark (*Carcharhinus limbatus*), tiger shark (*Galeocerdo cuvier*), and bull shark (*Carcharhinus leucas*). In Colombia, 60% of respondents reported that juvenile sharks were caught incidentally while fishing in coastal areas and 30% reported that mangroves are important for juvenile sharks. In Panama, 64% of respondents mentioned that juvenile sharks are common in landings and 60% highlighted mangroves as important nursery and feeding areas for sharks. In Costa Rica, 30% of respondents said juveniles were often caught with gillnets and 50% mentioned a sharp decline in shark abundance in the last decade.

In Colombia, Panama and Costa Rica, 96%, 85% and 95% of respondents respectively said that overall catch volume had declined compared to when they had started fishing. When asked about the reasons for the reduction, 90% of respondents in Colombia, 59% of respondents in Panama, and 89% of respondents in Costa Rica attributed it to overfishing. More specifically, the use of gillnets was highlighted as the main cause of decline in Colombia, whereas the increased number of fishers was the main cause perceived by Panama respondents. When asked about potential solutions to increasing fishing pressure, there was a generally positive reaction to responsible fishing zones. In Colombia, 98% of respondents stated that responsible fishing zones and MPAs generate significant increases in fish abundance, but mentioned that these areas should be established with scientific information and socioeconomic studies. In Panama and Costa Rica, 64% and 50% of respondents supported responsible fishing zones as a viable solution.

Across the region respondents believed mangroves are important, with 100% of respondents in Costa Rica, 92% in Colombia and 81% in Panama, highlighting the great value of mangroves not only for their lives and livelihoods but specifically as nursery habitat for commercially important fishery species, including sharks. However, respondents also observed declines in mangrove health and cited various reasons. The most important factor affecting mangroves in Colombia was general pollution (fuel, oil, plastic) with 33% of respondents citing it. In Panama, 43% of respondents mentioned agricultural fertilizers as the most damaging factor for mangroves, and in Costa Rica, 50% of respondents mentioned logging and pollution.

## Mobulid rays

Interviews revealed that the occurrence of mobulid rays (mantas and devil rays) in coastal areas is sporadic and limited to islands and seamounts. Manta aggregations peaked between December – April in Costa Rica and Panama. Away from these areas, fishermen state they observe them only rarely when they jump, yet confusion is common throughout the region as most rays (even demersal stingrays) are named “manta” or just “ray”, so Pacific cownose rays (*Rhinoptera steindachneri*) are commonly misidentified as mobulids by fishermen. Mobulid rays are caught incidentally in long line and tangle net fisheries. Most are discarded or utilised as bait in the long line fishery. The most commonly recognised species of mobulid from ID cards was *Mobula thurstoni*, the smoothtail devil ray/mobula or lesser devil ray. Giant mantas occur occasionally, but the incidental catch in long line fishery is rare (∼1-2 per year). Fishers interviewed in Colombia stated that “very large mantas’’ are frequently sighted off the coast of the Northern Choco but specific locations and timings were not obtained. A much greater threat to mobulid rays are gill nets, but fishermen state they actively try to avoid areas frequented by mantas as they can drag the net or boat dangerously, and cause damage to nets, boats and propellers.

Dive shop surveys in Panama and Costa Rica revealed that there are aggregations of reef (*Manta alfredi*) and giant mantas (*Manta birostris*) at specific sites. In Panama, manta rays are most frequently sighted by divers in Coiba National Park around March. In Costa Rica, manta ray occurrence was reported to peak between December and April in the southern part of Isla Tortuga and Punta Roca, Reserva Curú and Islas Negritos.

### Community site mapping

Interview respondents highlighted historically important areas for sharks and often recounted stories of particularly large sharks or pregnant females that were captured in shallow waters. Most critical habitats identified were associated with gulfs, mangroves, estuaries and deep sloping islands close to shore. In our analysis, we produced a map of critical habitat for sharks in the ETP according to fisher’s traditional ecological knowledge, ground-truthing and using small-scale fisheries landings data (figure 2).

**Figure 2.**
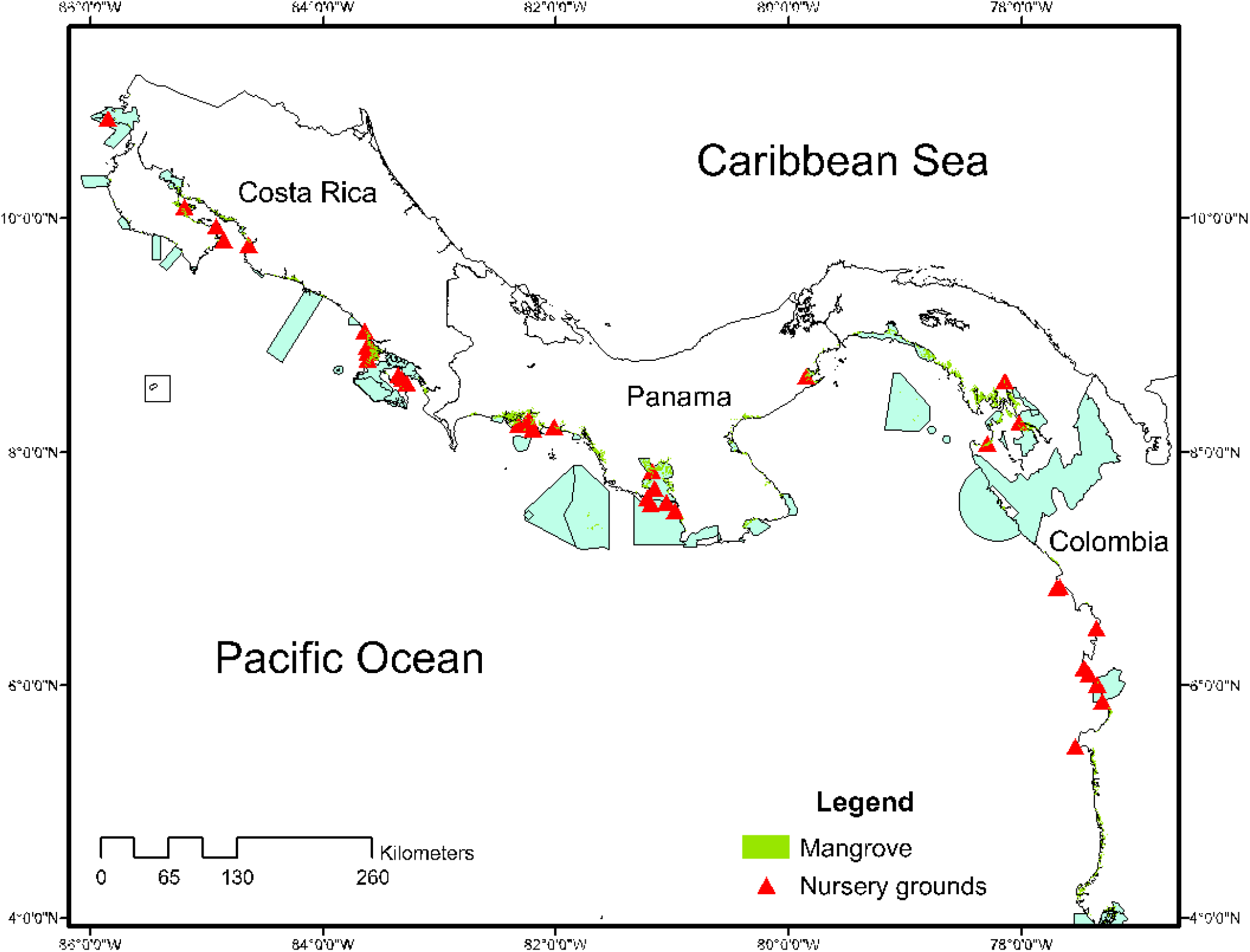
A map of anecdotal and observed locations of nursery areas for juvenile sharks in the Eastern Tropical Pacific, compared to existing networks of marine protected areas (blue shaded polygons).

### Small-scale fisheries landings

#### Colombia

In the northern Chocó region, the most frequently landed species in small-scale fisheries were *Sphyrna lewini, S. corona, S. tiburo*, and *Mustelus lunulatus* (Figure 3, 4 and Appendix 5), which is corroborated by a similar study (Díaz, Melo and Roa, 2017). The low mean weights for *S. lewini* imply that the fishery is dominated by juvenile landings (Figure 5). This region is divided by management regimes: The northern two-thirds of the coast is designated as an Exclusive Zone of Artisanal Fishing (ZEPA) that prohibits gill nets (Figure 1). The lower third is an area that, during the time this study took place, was open to almost all types of artisanal fishing as well as industrial shrimp trawling. It has subsequently been gazetted as a special management area called the Regional Integrated Management District (DRMI) of the Gulf of Tribuga, Cabo Corrientes. Biogeographically, the two areas are somewhat different in that the lower Gulf of Tribuga contains more extensive mangrove cover than the north. Landings analysis using *Index of Relative Importance* highlighted the correlation between species and these mangrove habitats (Figure 5) and also allowed for some evaluation of the effects of management on catch composition, such as a Utría National Park and the ZEPA.

**Figure 3:**
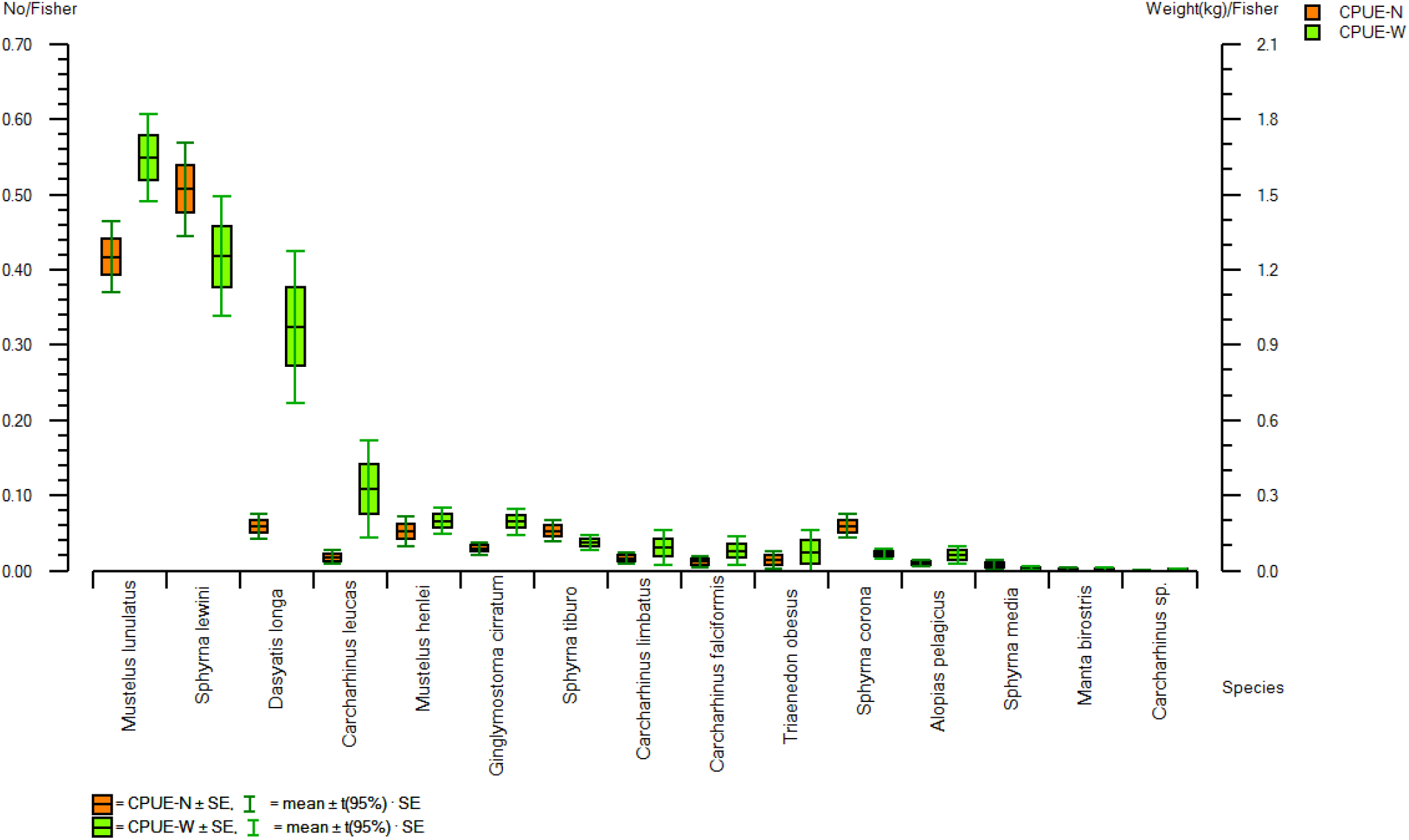
Catch per unit effort by weight (kg/fisher*day) in orange, and by the number of fish (N/fisher*day) in green, for all elasmobranch species recorded in landings from Northern Chocó, Colombia between 2010-2013

**Figure 4.**
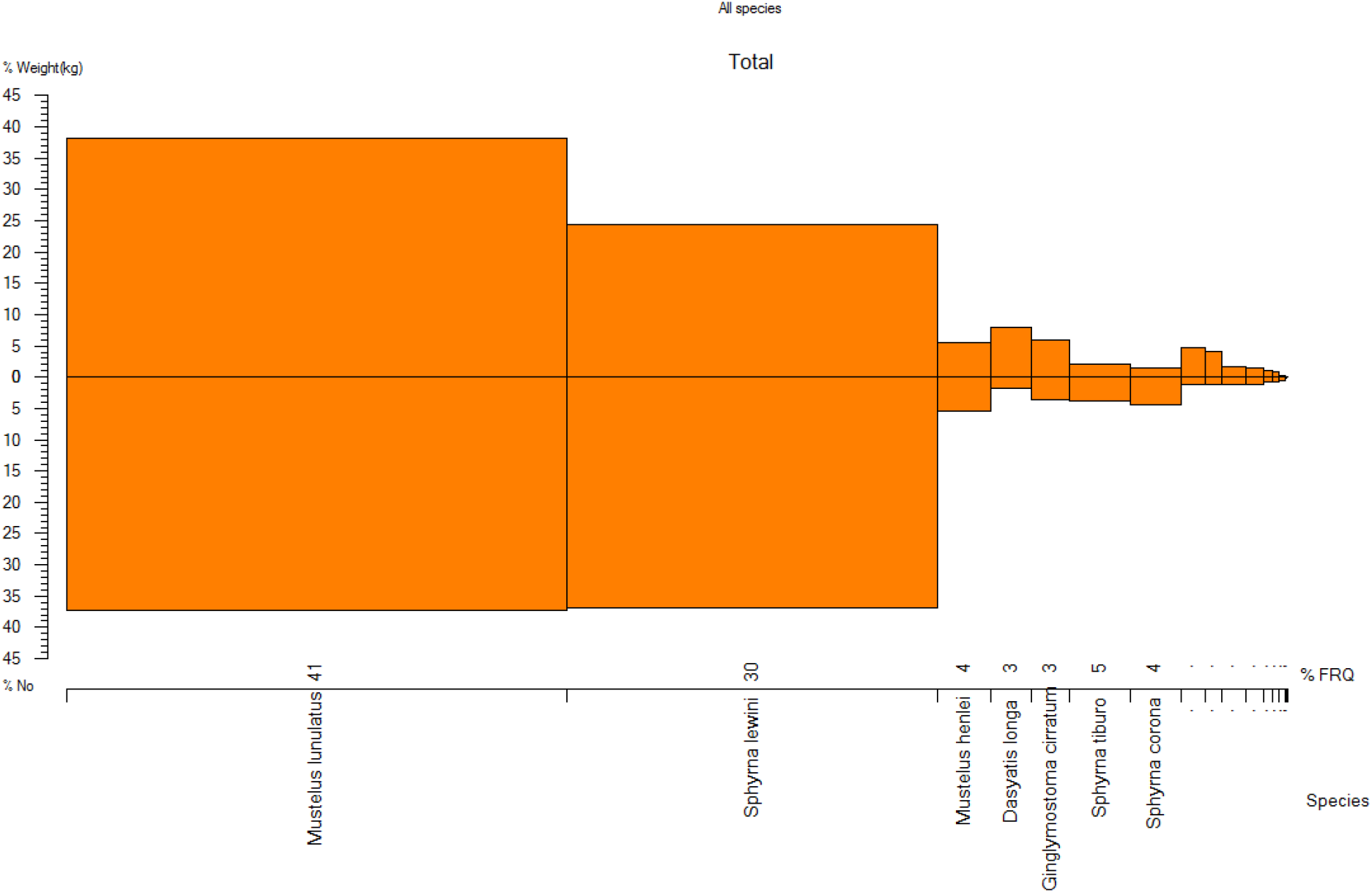
Index of Relative Importance for the top 7 elasmobranch species landed across all sites in the Northern Choco, Colombia between 2010 and 2013.

**Figure 5:**
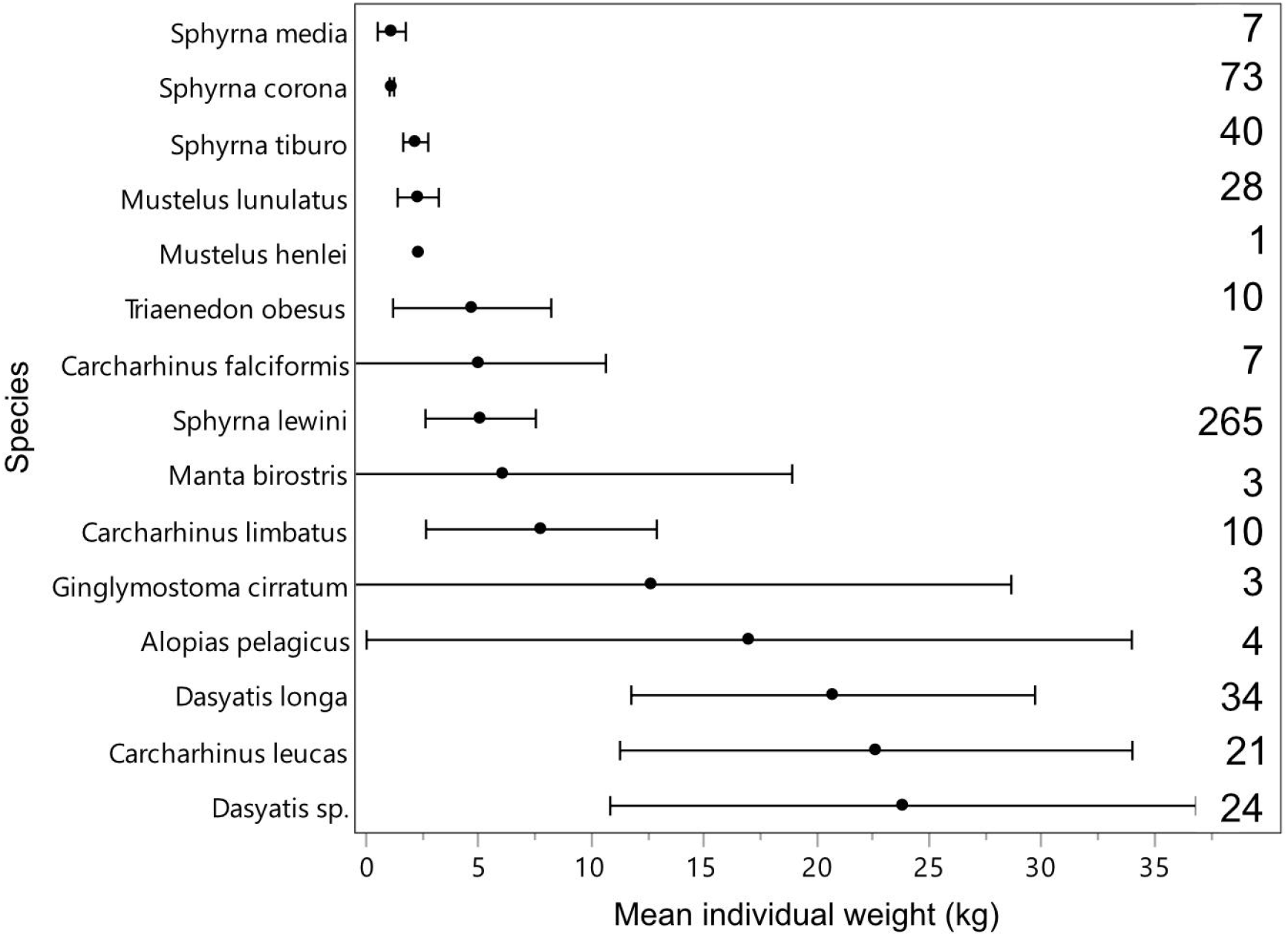
Mean individual weight in kilograms by species landed in artisanal fisheries of the Northern Choco, Colombia (total landed weight of individuals divided by the number of individuals landed). Error bars reflect 95% confidence intervals and numbers down the right-hand side reflect the sample size of each species.

The catch composition of elasmobranch species was similar throughout the region, and the management regime in place made no discernable difference to shark and ray abundance. In the Gulf of Tribuga in Colombia, where no regulations were in place at the time of data collection and where mangroves are abundant, showed higher mean CPUE for sharks and rays than the ZEPA across all gear types.

For *S. lewini*, peaks in relative abundance were seen from April-July (Figure 7). *Sphyrna lewini* was recorded in landings from 12 of the 17 locations. The mean individual weight of 2.58 kg suggests that the majority were juveniles (figure 5). Landings from the Gulf of Tribuga showed much higher CPUE than in the ZEPA, which one would expect given the greater mangrove habitat available, and prevalent use of gillnets in this area. Correspondingly, the ZEPA showed much higher weight landed by long lines and hand lines.

**Figure 6.**
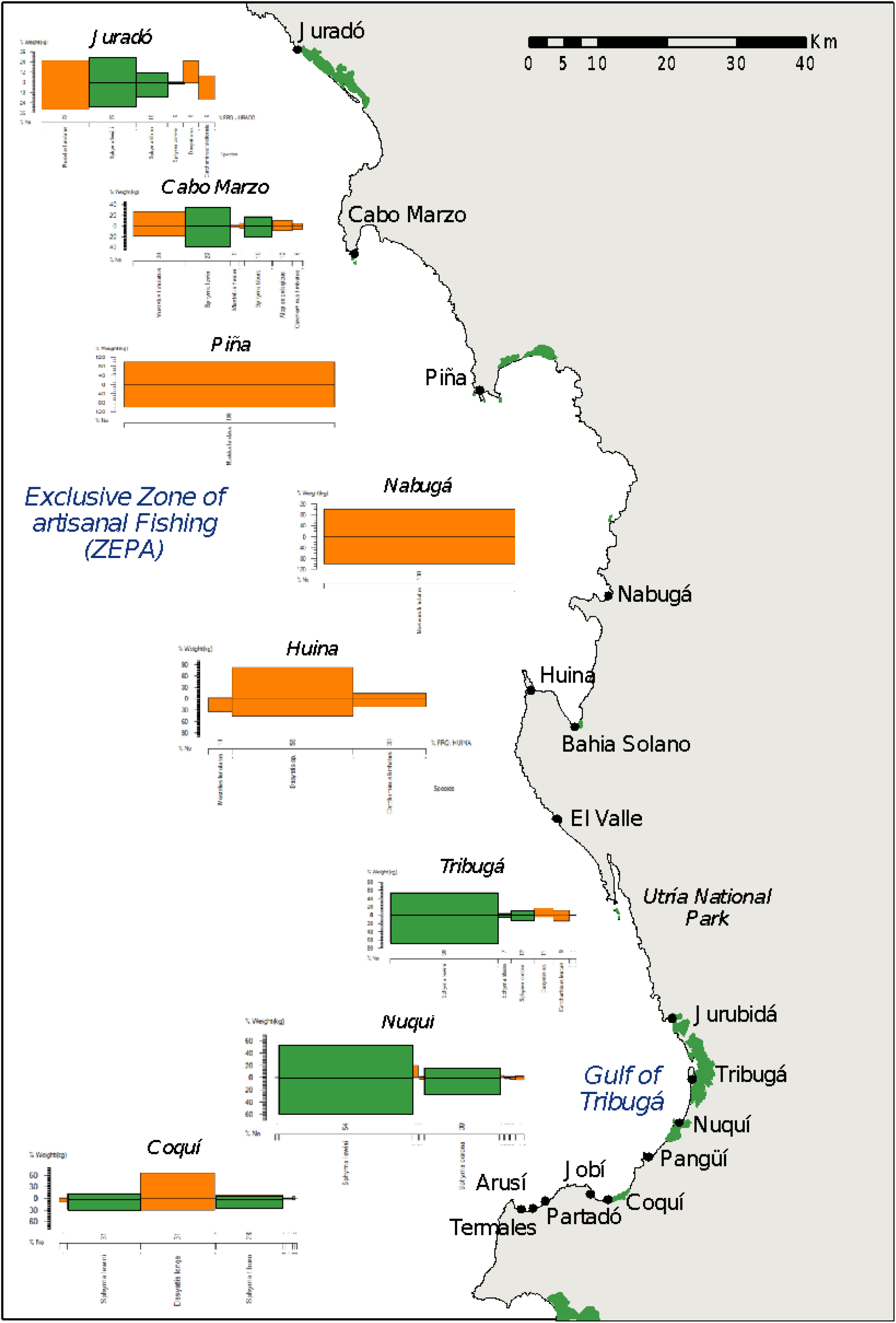
Map of Northern Chocó, Colombia showing landing sites and selected IRI analyses for species of sharks and rays. The size of the box represents a scale of importance as a proportion of the total catch by location. Hammerhead sharks (*Sphyrna* spp.) are shaded green. All other sharks and rays are coloured orange.

**Figure 7:**
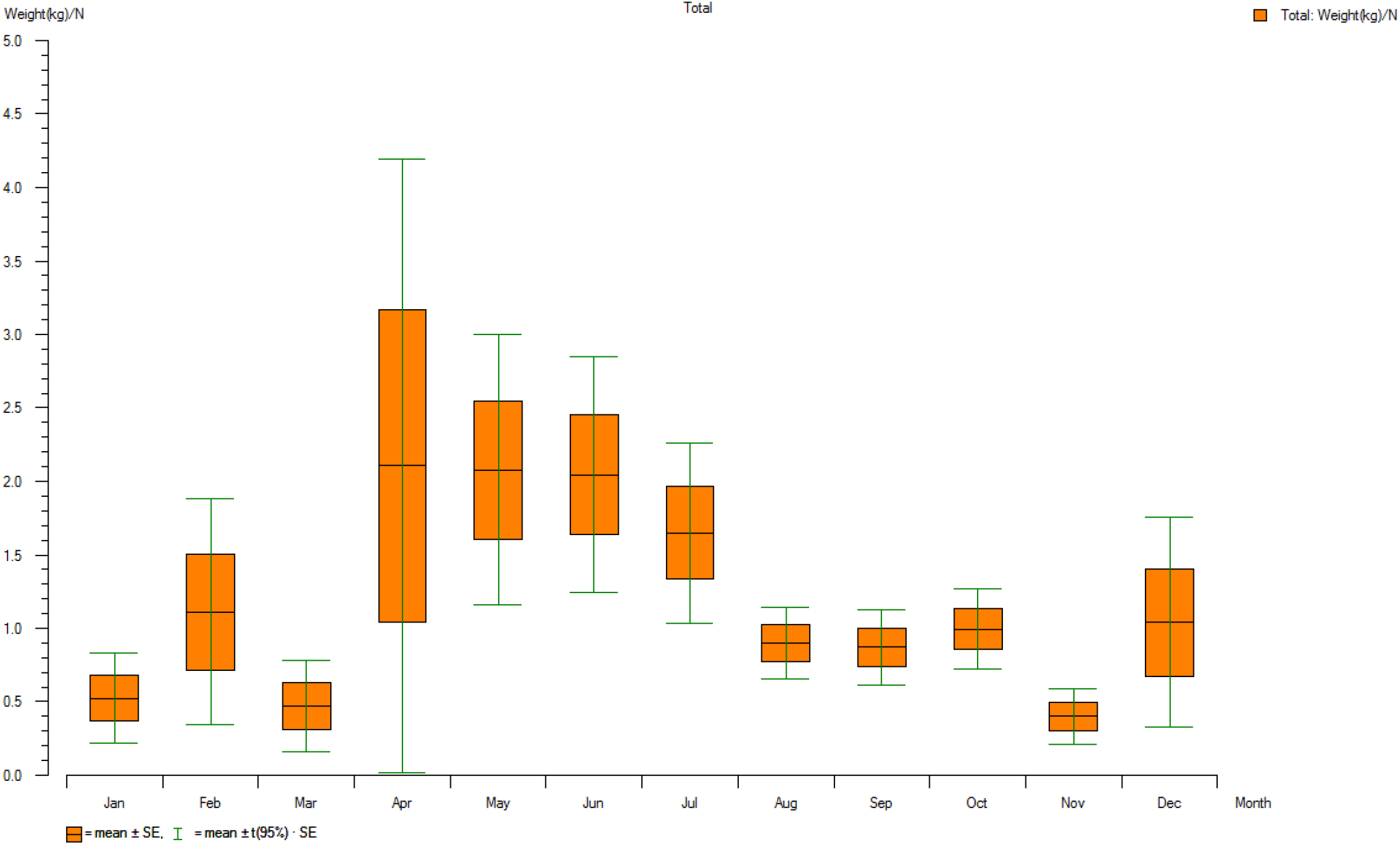
Mean CPUE (kg/fisher*day) of *Sphyrna lewini* by month in the Northern Chocó, Colombia.

#### Panama

Landings data from Panama were limited to 2 regions, the Gulf of Montijo 150 km west of Panama City, and the Gulf of Chiriquí in the west of the country. Individuals recorded in landings data were only identified to genus. Hence, Sphyrnid sharks (hammerheads) are likely to be representatives from all 5 locally occurring species. All other sharks were misidentified at the time of recording as *Carcharhinus porosus* (which only occurs in the Atlantic). The most commonly recorded species from the region was the blacktip shark, *Carcharhinus limbatus*. It is likely that multiple Carcharhinid shark species have been captured in these fisheries but are misidentified or only identified to genus (*Carcharhinus* sp.). No mobulid rays (grouped locally into “manta ray”) were recorded as landed in Panama records.

Although few records existed with both weight and number of individuals landed (N = 17), those for which the number was recorded were very young juveniles. The mean size of Sphyrnid sharks was 1.00 kg (from Cebaco, Hicaco, Santa Catalina sites) and Carcharhinid sharks being 1.94 kg (from Bahia Honda, Hicaco, Santa Catalina). The most important gear type by total cumulative landings was gill nets, but when coarsely standardised by effort (trip), gill net CPUE of sharks was lower than hand lines and long lines (Table 2).

**Table 2.** Combined total landed weight and catch per unit effort (CPUE) of sharks by gear type in the Gulfs of Montijo and Chiriquí, Panama.

| Gear Type | Total landings (kg) | N | Mean catch per trip (kg) | SE |
| --- | --- | --- | --- | --- |
| Hand line | 180.89 | 4 | 45.22 | 18.44 |
| Long line | 192.62 | 8 | 24.08 | 11.41 |
| Gill net | 1598.97 | 120 | 13.32 | 1.59 |

The majority of records came from the area of the Gulf of Montijo, with sharks caught throughout the year. Peaks in landings were seen in April & May and July & August (Figure 8).

**Figure 8.**
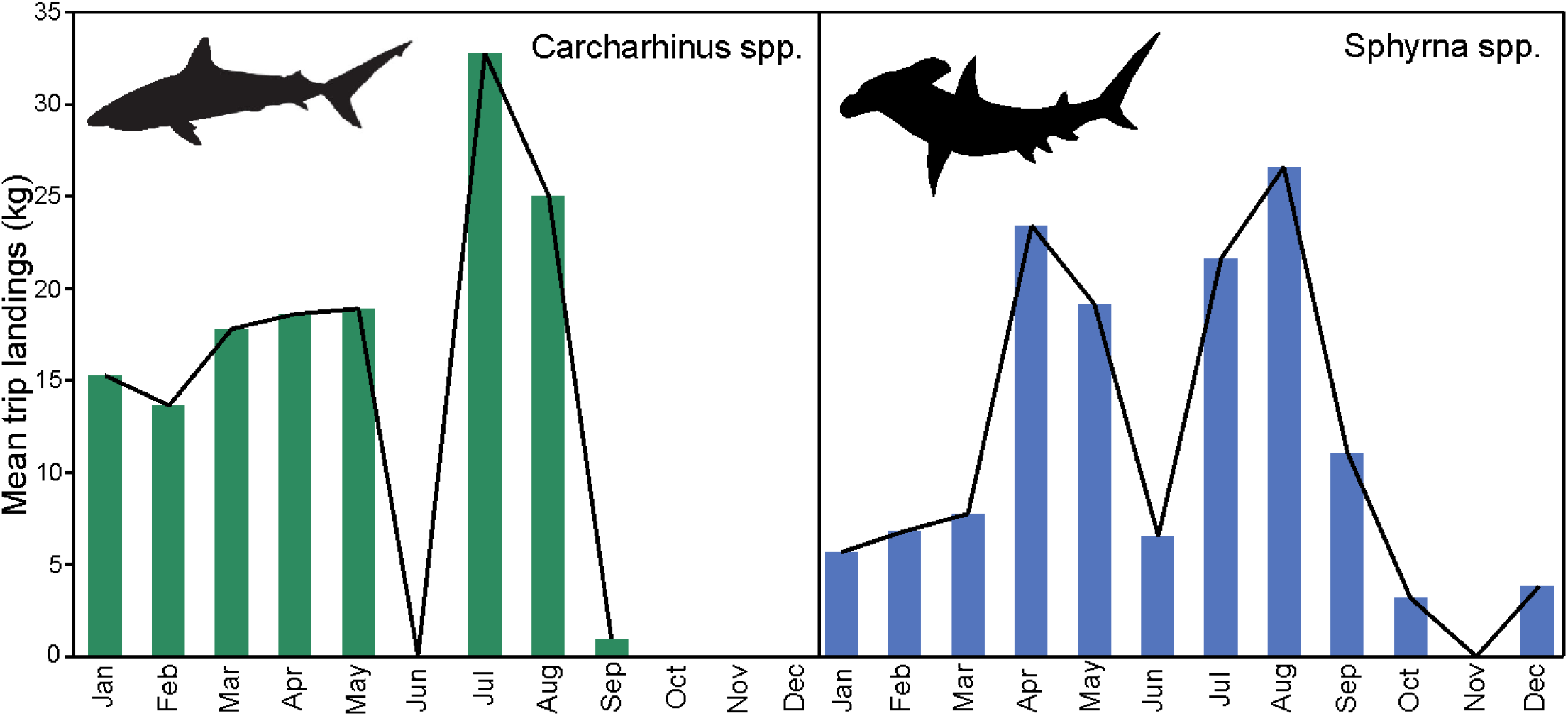
Mean catch per sample (trip) grouped per calendar month for *Carcharhinus* spp. and *Sphyrna* spp. in Panama between 2012-2014. Black continuous line added for visualisation purposes only.

### Inclusion of critical habitats in MPA network

From the 41 locations identified in this report as critical habitats for coastal sharks, 37% already fall within the protected area networks of the ETP (figure 2). For Costa Rica, 30% of the critical habitats identified in this study were inside marine reserves. This figure was 48% for Panama and 30% for Colombia. Our analysis of mangrove representation in the MPA network showed that Costa Rica has the best representation by proportion protected, followed by Panama (Table 3). Total mangrove area in Colombia is slightly higher than Panama, but needs better protection, as currently, only one-quarter of Colombia’s Pacific mangrove forests are under some form of protection.

**Table 3.** Representation of mangrove habitat in the Marine Protected Area Network of the ETP.

| Country | Total mangrove area, Pacific coast (km<sup>2</sup>) | Protected mangrove area, Pacific coast (km<sup>2</sup>) | % of mangrove protected |
| --- | --- | --- | --- |
| Colombia | 1506.1 | 418.3 | 27.7 |
| Panama | 1429.7 | 615.9 | 43.1 |
| Costa Rica | 390.3 | 229.0 | 58.6 |

## Discussion

Our study results provide evidence of the widespread usage of nearshore areas of the eastern tropical Pacific by sharks and maps these critical areas as an important baseline for future research and prioritisation of fisheries management in the region. The importance of mangroves for fisheries biomass is now well-established (Carrasquilla-Henao and Juanes, 2017), and the correlation between juvenile shark landings and the presence of mangrove forests was evident from interviews and catch records. Unsurprisingly, given the history of human association with mangrove forest resources in the region (López-Angarita *et al*., 2016) the majority of interview respondents were cognizant of the importance of mangroves as a resource base for their livelihoods and the life cycle of shark species. In our interviews, respondents recognised the importance of mangroves and river mouths as nurseries and pupping grounds, with some showing pictures of large pregnant sharks caught in the past. A large number of neonate hammerhead and blacktip shark landings were observed in fisheries from within and beside important mangrove forests where fishing is prohibited (Image 2).

**Image 2.**
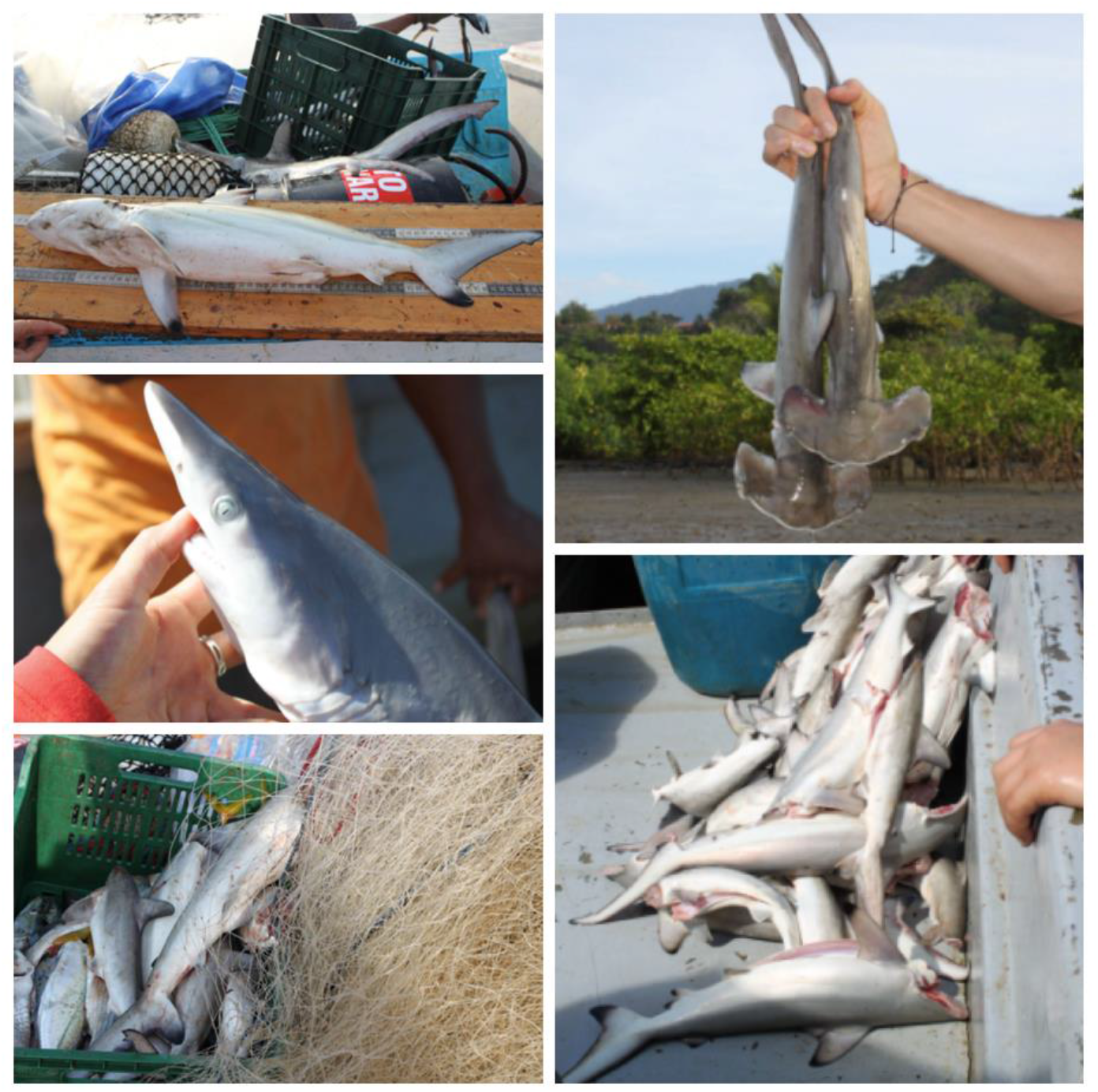
Neonate sharks observed being landed in a Costa Rica gill net fishery. Most individuals were scalloped hammerhead (*Sphyrna lewini*) and blacktip sharks (*Carcharhinus limbatus*). © Talking Oceans.

Most large mangrove systems in the region are offered some level of protection from clearing and cutting, though human impacts vary enormously by location (López-Angarita, Tilley, Hawkins, *et al*., 2018). Costa Rica has well-established policies dating since 1998 that protect mangrove forests (López-Angarita *et al*., 2016), and is the country with the highest proportion of mangrove forests inside reserves. Our results suggest that 63% of identified critical habitat spots across all countries are not included inside protected areas. Furthermore, those areas that are under protection still seem to be actively fished by surrounding communities. Fishers appeared to be aware of fishing restrictions in these areas, but compliance varied widely according to location.

Variability of volume and frequency of sharks in catches throughout the year suggests there are important seasonality considerations in managing coastal fisheries for shark conservation. However, the lack of fisheries independent data and sufficiently standardised sampling of landings precludes us from drawing strong conclusions regarding the drivers of abundance across seasons. In other areas, reproductive events, and movement to inshore areas in search of seasonal prey items (e.g. sardines) have been shown to increase the numbers of sharks in coastal waters (e.g. (Chapman *et al*., 2009; Speed *et al*., 2010; Bangley *et al*., 2018; Zanella, López-Garro and Cure, 2019)). Mobulid rays appear to have very limited usage of coastal habitats, occurring only at aggregation spots of upwelling around islands or seamounts. A number of these sites exist within the ETP, and hence are a habitat type in addition to mangroves that should be considered for greater protection.

The scalloped hammerhead, *Sphyrna lewini*, was the most common shark species in landings across the areas surveyed. Interview respondents identified areas where juveniles are often caught, and many of them were located in or near mangrove forests, which was corroborated by our spatial analysis and follows findings of Díaz et al. (2017). The low mean individual weight of landings at less than 3 kg suggests that juvenile sharks are being landed in high frequencies, and these catches of small individuals were associated with fishing trips using gill nets. Interview responses suggest that there is a general perception of a decline in fisheries productivity in Colombia, Costa Rica and Panama and that overfishing is the leading cause through increasing effort and use of gill nets.

Throughout the region, in parallel with much of the developing world, the near-ubiquitous use of gill nets has had negative consequences on large-bodied, nearshore species (Dulvy *et al*., 2014). Local fisheries stakeholders are well aware of the damage gill nets cause, but gill nets are often perceived as the only effective gear in subsistence fisheries specifically, probably because gill nets continue to provide economically viable catch rates at dangerously high levels of effort (Northridge, 1991). Recent studies showed that fishers consider gear regulations, such as prohibiting small mesh gill nets of particular importance to the recovery of fish stocks (Saavedra-Díaz, Pomeroy and Rosenberg, 2016), and without new measures to manage fisheries, local fishers risk further descent into a social-ecological trap, where competition for a dwindling resource drives increased poverty among resource users (Cinner, 2011).

Shark conservation and fisheries management policies at local levels generally focus on restrictions of trade, fishing gear or fishing area (Shiffman and Hammerschlag, 2016). As to be expected, there is evidence that banning gillnets can result in the recovery of species susceptible to gill nets (Pondella and Allen, 2008). The socio-economic vulnerability of fishing communities to management restrictions is not equal (Tilley *et al*., 2018), but where gillnets are banned in the ETP (ZEPA, Choco, Colombia and Isla Chira, Costa Rica), fishers’ testimonies were positive, stating that abundance of fish had increased and claimed that fishing was more profitable than before. A general lack of control of fishing effort is common to small-scale fisheries in Latin America (Salas *et al*., 2007) but their informal and often subsistence nature makes entry restrictions impractical (Pomeroy, 2012).

Despite evidence suggesting that top-down fisheries management policies have not provided sufficient protection for sharks in the ETP (Brodzinsky, 2011), the establishment of responsible fishing zones over the past decade has shown demonstrable ecological gains (Vieira, Diaz and Díaz, 2016; López-Angarita, Tilley, Díaz, *et al*., 2018), and social gains as a model of co-management. In Costa Rica’s Gulf of Nicoya, fishers claimed that sharks, turtles and dolphins have been seen returning during the shrimp closed season (when gillnets are more strictly prohibited). Area closures can protect adult sharks through some of their activity space (Knip, Heupel and Simpfendorfer, 2012) or safeguard gravid sharks (O’Keefe, Cadrin and Stokesbury, 2013), but are most suitable for nursery ground conservation (Heupel, Carlson and Simpfendorfer, 2007).

Responsible fishing zones are a form of co-management of resources and allow for the development of social capital in Latin America (Salas *et al*., 2007). In general, where responsible fishing zones had been established, fishers said they were important and that catch had increased, but research in Colombia suggests that the ZEPA may not have reduced overall effort, but rather merely displace it to other areas (López-Angarita, Tilley, Díaz, *et al*., 2018). Analysis of landings and catch per unit effort support the anecdotal success attributed to responsible fishing areas in terms of fisheries productivity (López-Angarita et al., 2018), but more studies are needed to assess their protection for sharks in particular. News of these successes has spread to other areas in Colombia and Costa Rica with newly established areas such as the Regional District of Integrated Management (DRMI) in the Gulf of Tribuga in 2015 (Satizábal, 2018). This area promotes the conservation and sustainable use of biodiversity, and certain habitats are zoned for conservation, controlled use and extraction (Rojas et al., 2019). In Panama, interviewed fishers were fearful of restrictions of gill nets because it is their main fishing method, and only recently in 2019 has the first co-managed responsible fishing area been established (ARAP, 2019).

## Conclusions and recommendations

Our results suggest that responsible fishing areas are working to foster recovery of coastal fish stocks and should be more broadly adopted by small-scale fisheries in the region. The use of interviews to record ecological knowledge was particularly pertinent in the ETP where information is scarce or lacking. Interviews highlighted that, in general, the political will exists in fishing communities to protect natural resources for traditional livelihoods and human wellbeing. Recording fisher knowledge and aspirations as part of the management process serves to legitimise their role as stewards of marine resources and increases the likelihood of sustainable resource use (Pomeroy *et al*., 2015; Tilley *et al*., 2019).To capitalise on this, local governments must consider inclusive governance approaches such as the development of co-managed responsible fishing zones. These initiatives will need to establish locally legitimate, co-generated monitoring programs to collect and utilise data to drive science-based decision-making for the sustained recovery of local stocks (Hilborn *et al*., 2020). This study highlights the importance of traditional ecological knowledge in identifying critical habitats and prioritising research of endangered species.

## Footnotes

1 Inter-American Tropical Tuna Commission, 2011; *Costa Rica*, General Assembly Law 8436-2005; *Ecuador*, Ministry of Agriculture, Livestock, Aquaculture and Fisheries (MAGAP) agreement 116-2013; *Colombia*, Ministry of Agriculture and rural development, Resolution 434-2019

2 IPOA-Sharks, FAO, http://www.fao.org/ipoa-sharks/en/

